# Functional Roles of the World’s Recovered Predators

**DOI:** 10.64898/2026.01.14.699583

**Authors:** Ishana Shukla, Julia D. Monk, Justine A. Smith

**Affiliations:** Department of Wildlife, Fish, and Conservation Biology, University of California, Davis, CA, USA 95618; Department of Environmental Studies, New York University, New York, NY, USA 10003

**Keywords:** Restoration, predator, conservation, community ecology, reintroduction, rewilding

## Abstract

Terrestrial predator populations have declined globally due to anthropogenic pressures with cascading consequences throughout ecosystems. As a result, recent efforts have sought to restore predator populations both passively (via habitat conservation and connectivity) and actively (via conservation translocations and reintroductions). Predator recovery efforts often cite restoration of ecological responses as a central goal, yet few studies monitor and report ecological outcomes of predator restoration. We conducted a comprehensive global meta-analysis of ecological responses to predator recovery, specifically investigating drivers of the magnitude of responses and, crucially, their alignment with local conservation objectives. Our analysis revealed that recovered predators overall elicit strong responses from the wildlife community, with most responses aligning with local conservation management preferences. Recovery effort techniques and biological community characteristics influenced the strength of community response and alignment with conservation goals. Although our results suggest that predator recovery is largely beneficial to ecosystem function, clarifying the goals of predator restoration will be essential for successful evaluation of ecological restoration and development of coexistence strategies.

## Introduction

In an era defined by global defaunation and species declines, the indirect effects of species loss on ecosystem function are increasingly apparent. The extirpation of terrestrial large predators in particular have been implicated in subsequent cascading impacts on biodiversity, productivity, disease, and nutrient cycling (Estes et al. 2011, Ripple et al. 2014, Atkins et al. 2019). Recognition of these impacts has motivated efforts to facilitate the restoration of extirpated predators into their historic ranges via active reintroductions (Armstrong and Seddon 2008, Resende et al. 2020, Thomas et al. 2023, Serota et al. 2023). In addition to these active restoration programs, conservation actions that prioritize connectivity or landscape restoration have facilitated passive predator recovery as animals take advantage of improved habitat quality or reduced threats, dispersing into their former ranges (Nicholson et al. 2014, Buderman et al. 2018, Engebretsen et al. 2021). Predator restoration actions often focus on two primary goals: the recovery of the predator population itself and the restoration of vital ecosystem functions (Wilmers et al. 2025). Efforts focused on improving the conservation status of predators have shown marginal success in both survival and reproduction metrics (Thomas et al. 2023). However, despite garnering widespread attention and, at times, controversy, the effectiveness of predator recovery projects at eliciting desired community outcomes and reinvigorating targeted ecological processes has not been evaluated at scale (Ritchie et al. 2012, Alston et al. 2019). Though specific case studies have been hotly debated (perhaps most notably the question of wolf-induced trophic cascades following active reintroductions in the Greater Yellowstone Ecosystem; e.g Ripple and Beschta 2003, Kauffman et al. 2010, Marshall et al. 2013, Brice et al. 2022, 2025, Wilmers et al. 2025), the broader scientific question of how environmental and management parameters mediate the strength and predictability of the ecological outcomes of predator restoration is prime for more thorough investigation.

Community responses to restored predators can vary widely, encompassing everything from direct changes in prey abundance and behavior to cascading effects on lower trophic levels and the abiotic environment (Estes et al. 1978, Terborgh et al. 2001, Ripple and Beschta 2012, Stier et al. 2016, Clark et al. 2016). It stands to reason that the method of predator recovery (passive vs. active) could impact the nature of species interactions and, consequently, the magnitudes of ecological responses (Fig. 1), though this hypothesis has not been widely investigated. Passive recoveries occur when a predator naturally disperses into and recolonizes an area in their former habitat range, or when a threat is mitigated (e.g. decreased hunting quotas, increased habitat conservation) and predator populations rise to their historic levels (Ewen et al. 2012). In contrast, active recoveries, which refer to translocations and captive releases, involve the intentional movement of animals from one location to another by humans (Fischer and Lindenmayer 2000, Ewen et al. 2012). Translocations, captive releases, and dispersals may produce differing levels of establishment success and operate at different timescales, thus impacting the magnitude of effect that the predator elicits from the community (Fischer and Lindenmayer 2000, Jule et al. 2008, Resende et al. 2020); Fig. 1). Active recoveries may promote higher rates of reestablishment as they are carefully planned before predator reintroduction and often have elements of sustained active management. However, introducing naive predators could offset these advantages as their sudden introduction into a new system could lead to disadvantages in hunting success and competition, thus decreasing their impacts on the community (Resende et al. 2020). Passive recoveries gain the advantage of a slow reintroduction, wherein the recovering predator population can gradually accumulate place-based knowledge as they continue to disperse and expand their ranges. However, in passive recoveries, the stressor that caused the initial predator extirpation may still be present, which then only serves to re-expose the unequipped predator population to the disturbance (Palmer et al. 2010) and may thus limit ecological effects.

**Figure 1.**
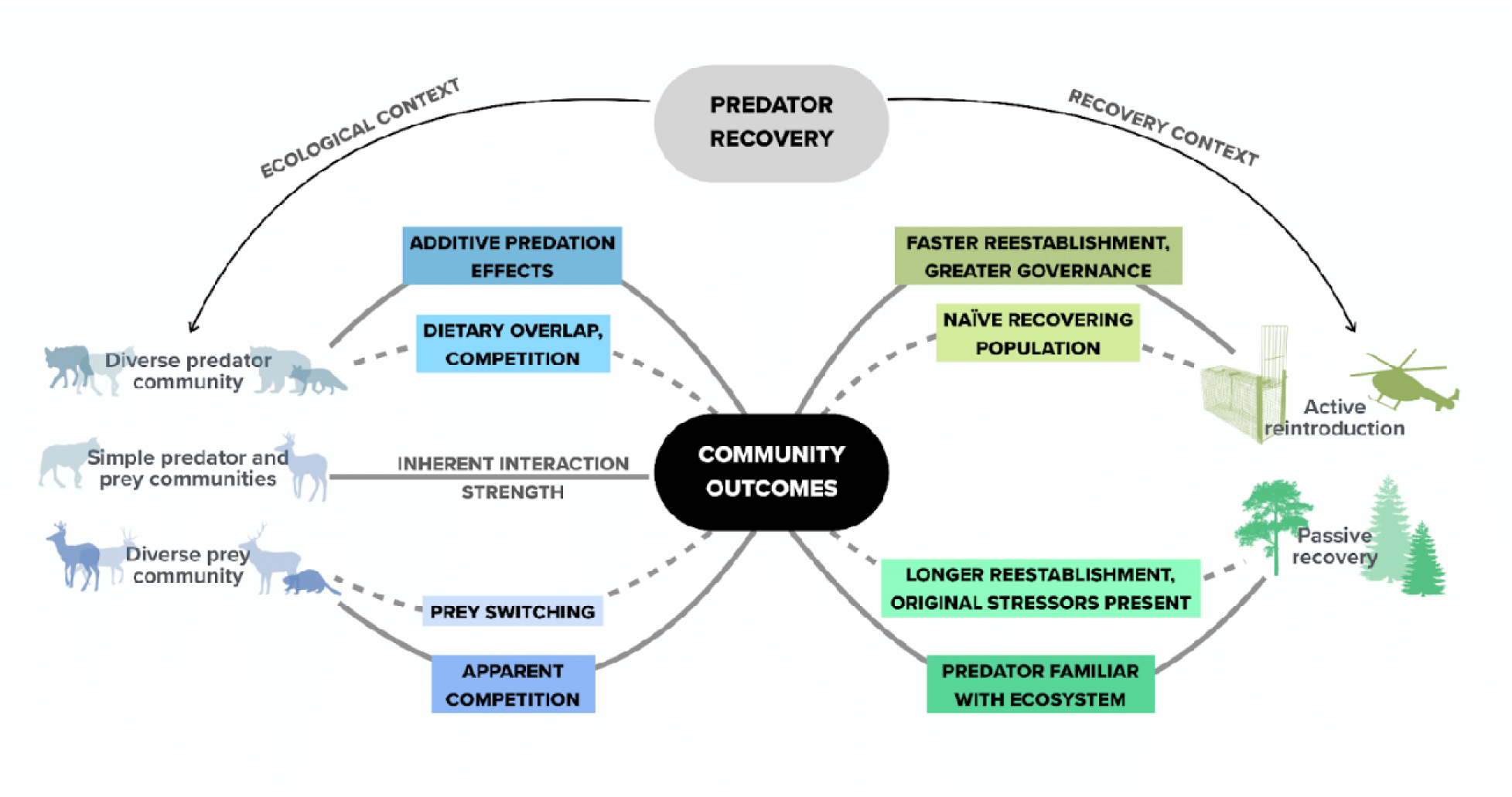
Factors influencing the magnitude of community responses to a recovered predator. Solid lines indicate interactions that could increase the magnitude of community outcomes, while dashed lines indicate interactions that could decrease the magnitude.

The pre-existing structure of the biological community is also likely to have substantial impacts on the community outcomes of a recovering predator (Samhouri et al. 2017). The diversity of the existing prey community might influence the extent that a predator can impact a community, as multiple prey species could allow for either prey switching behavior (reducing predator impact; Garrott et al. 2007) or cause apparent competition (increasing predator impact; Holt and Bonsall 2017). Likewise, existing predator diversity is also likely to affect the magnitude and breadth of community outcomes. If a recovering predator species’ diet largely overlaps with an extant predator population’s, the functional redundancy could reduce the overall impact on the community (Rosenfeld 2002). Conversely, additive predation effects could increase the impacts on prey (Roemer et al. 2001). Overall, community ecology theory suggests that simple systems (i.e. single predator species and single prey species) would innately contain higher levels of interaction strength between the species, and thus produce a stronger community response than in more biodiverse and complex communities (Kokkoris et al. 2002, Berlow et al. 2004, Monk and Schmitz 2022).

Beyond ecological outcomes, little evidence exists on how recovered predators interact with adjacent human populations, or whether observed ecological and social outcomes map onto initial conservation goals. Recoveries of large predators in particular pose unique socioecological challenges due to their extensive habitat needs, long life histories, and higher risk of human-wildlife conflict (Stier et al. 2016, Hill et al. 2019). Many recovered predators exist in shared and mixed-use landscapes, increasing the chances of contact with humans. While some outcomes of human-predator interactions are positive (e.g., invasive pest control), large predators are more often implicated in negative outcomes such as livestock depredation or direct risk to human safety (Harihar et al. 2011, Twining et al. 2021). If nearby human communities are not receptive to the recovery of a large predator population, the establishment of the population may be slowed (due to harassment, retaliatory killing, or illegal take), therefore reducing the subsequent impacts on other species in the community (Treves et al. 2017).

We conducted a global meta-analysis to investigate the environmental factors driving the nature and magnitude of community responses to predator recoveries and their alignment with conservation goals. Along with recovery mode (active vs. passive) and existing ecological community structure, we assessed myriad other ecological and social factors that potentially influence community responses and conservation alignment, such as predator characteristics (e.g., mass, trophic level), latitude, disturbance history (e.g., length of extirpation), and governing body that led the recovery. By uncovering the drivers of outcomes of predator recoveries, we offer insight into whether and when restored predator populations measurably impact ecosystem functioning, and whether these outcomes match the expectations and goals of those invested in predator recovery.

## Materials and Methods

### Data Collection

To assess drivers of recovered predator community outcomes, we conducted a systematic review. We searched Web of Science and Google Scholar with the search terms “Apex” OR “predator” OR “predat*” AND “rewild*” OR “reintrod*” OR “translocat*” OR “recovery” OR “restor*” OR “restoration” (Fig. S1). If the article’s title or abstract included one or more of our search terms, we then scanned the full publication to see if it met our eligibility criteria. To be eligible for inclusion in our meta-analysis, papers had to: i) focus on a recovered (either passive or active) terrestrial vertebrate carnivore (species whose diet is >50% animal matter); and ii) report empirical data on a quantitative response from the community, though those who did not report community responses were recorded for analysis of the global distribution of such predator recoveries (Fig. 2). We then performed both a forwards and backwards search by searching both the citations and references of each manuscript to further identify any other peer-reviewed articles on predator recovery outcomes. Finally, we located other reports of community responses to predator recoveries via the International Union for the Conservation of Nature’s (IUCN) Global Re-Introduction Perspectives Series (n = 7). Our literature search yielded 33 recovery projects with an effect size of 69 associated community outcomes..

**Figure 2.**
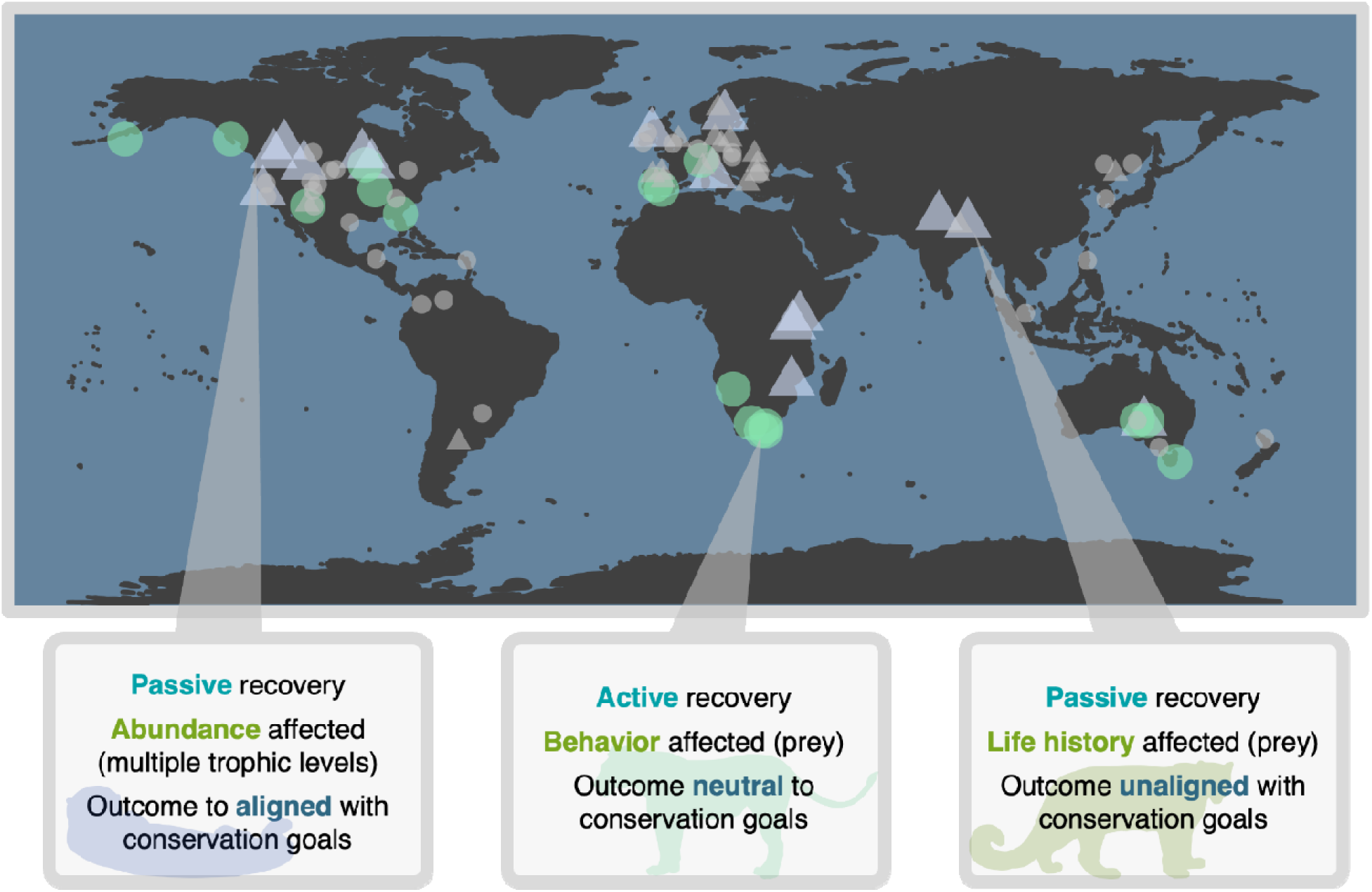
Passive (blue triangles) and active (green circles) predator recovery sites that have documented community responses included in the meta-analysis. Small, light grey shapes indicate existing studies on predator recovery for which published studies of community responses were not found. Examples from left to right: i) Passive sea otter (*Enhydra lutris*) recovery in the eastern North Pacific increased biomass of finfish and small invertebrates and decreased biomass of large benthic vertebrates (Gregr et al. 2020); ii) Prey became more diurnal in areas with reintroduced African lions (*Panthera leo*) and spotted hyenas (*Crocuta crocuta*) (Tambling et al. 2015); iii) Snow leopard (*Panthera uncia*) recovery reduced Himalayan tahr abundance (Lovari et al. 2009).

### Data Preparation

We compiled data on each species and associated community to explore the influence of ecological and recovery variables on the magnitude of community responses elicited by the recovered predator (Table S1). We recorded the taxonomic family of the predator, trophic level (apex or subordinate predator), predator mass, latitude of the recovery area, and existing wildlife community composition at the site of predator recovery (i.e., multiple vs single predator in the same trophic level, multiple vs single prey species in the recovered predator’s diet). To investigate the influence of recovery characteristics on community outcomes, we also extracted data on the time since the predator was extirpated from the site, the type of reintroduction (i.e., active or passive), the time since the initial recovery, the governing body that led the recovery (if any), and the GDP of the country where the recovery took place.

We also recorded various characteristics on the predator-induced community response. We first recorded the category of community outcome (i.e., changes in abundance [e.g., lower trophic level abundance or density], behavior [e.g. spatial or temporal activity], biogeochemical cycling [e.g., nitrogen cycling, bioturbation], life history traits [e.g., survival or reproductive parameters], or biodiversity [e.g. species richness]). We then recorded whether the outcome of the predator was aligned (i.e. invasive species control), unaligned (e.g., an increase in human-wildlife conflict), or neutral (e.g. shifts in activity patterns) with respect to local conservation goals or management preferences (Harihar et al. 2011, Ford et al. 2015, Twining et al. 2021). We sourced information regarding intention and alignment directly from the article. We finally recorded if active recovery projects reported a stated goal, and if the resulting recovery outcome was intended or unintended. Unintended consequences occurred if the main goal of the predator recovery was simply to bolster low populations (e.g., cheetahs *Acinonyx jubatus*), yet effects on other species in the community were documented, whereas intended outcomes occurred when a primary goal of the predator recovery was to reinstate the predator’s functional role in the community (e.g., sea otters *Enhydra lutris*; Gregr et al. 2020, Welch et al. 2022). We sourced information regarding intention and alignment directly from the article (Box 1).

#### Figure 5, Box 1. Towards Intentional Restoration: Evaluating Restoration Goals and Alignment with Conservation Values.

Intentional restoration of the functional roles of predators requires the establishment and evaluation of explicit conservation goals beyond predator recovery. However, some effects of predator restoration may not align with conservation goals (Fig. 5). Predator restorations with explicit intended ecological outcomes (e.g. control of invasive prey population) may produce consequences unaligned with conservation priorities (e.g. predation on endangered prey, Stepkovitch et al. 2023). Conversely, restorations with no intended ecological contribution may produce social or ecological outcomes that are aligned (e.g. regulation of mesopredators and release of their prey, Jiménez et al. 2019) or unaligned (e.g. conflict with livestock, Harihar et al. 2011) with conservation goals. Thus, while a crucial first step is to develop, monitor, and evaluate intended functional role restoration of predator recovery, broader monitoring of biological communities and engagement with local stakeholders is essential to build a comprehensive understanding of the indirect effects of restored predators. Without this more holistic approach, unintended effects are likely to be overlooked and underreported in the scientific literature, limiting opportunities to learn from previous restoration efforts and improve upon restoration practice.

In our comprehensive review, we found that a majority of ecological responses to predator restoration were unintended. Projects focused on predator recovery alone may thus miss opportunities to sufficiently evaluate cascading effects, develop conservation plans for native species, or prepare local communities for potential conflicts. Fortuitously, our review indicates that 77% of reported unintended effects are aligned with local conservation values. One of the reasons that unintended outcomes may often align with other restoration goals is that many conservation problems can stem from a similar underlying cause. For example, in a system in which a top predator has been lost and invasive prey have become overabundant, both problems can be ameliorated by predator recovery initiatives.

Despite most unintended consequences aligning with conservation goals, the other 23% require careful attention. Of the few cases in which predator outcomes were unaligned, many occurred in systems where many of the extant species are already at risk of extinction. In ecosystems that continue to experience severe anthropogenic threats, restoring predators may not reverse biodiversity declines or in some cases amplify them. Intentional predator introductions or translocations for biological control purposes have regularly resulted in non-target effects on native biota, particularly when prey animals are naive to predator cues and the predator is a diet generalist (Simberloff and Stiling 1996, Louda et al. 2003, Watari et al. 2008)). Such effects may be stronger in response to native predator restoration the longer the predator has been absent from the system (Berger et al. 2001), although prey in multi-predator systems are expected to better retain antipredator behavior (Blumstein 2006). For the field to progress towards better predictive evaluation of predator recovery programs, such complexities must be considered in advance and be paired with formal before-and-after monitoring of ecological and social responses.

The community responses themselves varied widely, and ranged from changes in stream morphology to prey behavior. To standardize between systems and outcome categories, we extracted the community metric when the predator was extirpated or at low density, as well as the community metric once the predator had been recovered. If the community metric was stated in terms of a gradient, we extracted the minimum and maximum value (Fig. S2). If numerical values were not provided in the text, we extracted the data from figures via WebPlotDigitizer (Rohatgi 2022). We then used a response ratio to compare community metrics between sites:

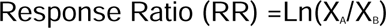

where X_A_ = the community metric in the extirpation (or reduced population) of the recovered predator, and X_B_ = the community outcome after the predator recovery. We then used the absolute value of response ratio to account for any cascading effects. If the study reported indirect responses across trophic levels (e.g. sea otter predation influencing the abundance of benthic invertebrates, cascading down to plankton abundance), we considered a change in each trophic level a separate predator outcome (Gregr et al. 2020).

### Assessing publication bias

We took a number of measures to address possible instances of publication bias. While conducting our literature search, we included gray literature, such as dissertations, theses and agency reports (n = 6). To further examine any potential bias in our completed dataset, we constructed a funnel plot and mapped the effect size residuals (response ratio) against the standard errors (Fig. S3). Finally, we employed an Egger’s test to measure any possible asymmetry (z = −0.3266, p = 0.74), and did not find any evidence of such (Egger et al. 1997).

### Statistical Analysis

To evaluate the recovery characteristics and ecological variables that influenced the magnitude of community responses to recovered predator populations, we evaluated a suite of candidate models. We constructed each candidate model as a mixed effects model with the *glmmTMB* package in R, each with a gamma distribution, recovery characteristics and ecological variables as fixed effects, and study and predator species as random intercepts (R Core Team 2018). We tested for multicollinearity with a variance inflation factor (VIF) test, and with all VIF values equaling approximately 1, we found an absence of collinearity. We then evaluated the best performing models and selected the top model based on the lowest AIC value (Symonds and Moussalli 2011) (Table S2).

To examine the factors driving whether these community outcomes were aligned, neutral, or unaligned in respect to local conservation and management preferences, we built a suite of ordinal logistic regression models with the *ordinal* package in R. We selected fixed effects from the same variables as our linear model competition and again included study and predator species as random intercepts. As our intended purpose was to investigate the driving factors of outcome alignment in relation to significant impacts from the human perspective, we only included studies in which community metrics in pre- and post-predator recovery were significantly different (n = 43 community outcomes).

## Results

Our final synthesis included data from 33 predator recoveries reporting 69 community outcomes, spanning 19 predator species from five taxonomic families (Fig. 2). Of the 112 predator reintroductions reported from 1976-2023, only approximately 30% of the projects monitored subsequent outcomes on other species in the community or ecosystem processes. The most represented predator families were *Felidae* (42%) and *Canidae (36%),* followed by *Mustelidae* (16%), and *Accipitridae* and *Dasyuridae* (2%, 2%). The mean recovered predator mass was 48 kg, and apex predators accounted for 88% of the cases with reported community outcomes. Most predator populations in our meta-analysis recovered passively (57%). The most commonly reported community outcomes were changes in lower trophic level abundance (50%), followed by changes in behavior (e.g, shifts in temporal activity) (37%), life history (e.g, reproductive rates) (5%), and finally biodiversity (e.g. lower trophic level richness), and biogeochemical cycling (e.g. changes to nutrient cycles), with each at 3%. Of the projects that had explicit recovery goals of lower-trophic level regulation (n = 19), only 35% of the resulting community outcomes were intended. However, of all community outcomes, 55% were aligned with local conservation or management preferences, 36% were neutral, and only 5% were unaligned.

The magnitude of community responses to predator recovery varied with the type of recovery, as well as the existing predator community composition. Our top-performing model accounted for ∼9% of the variation in community response. Aligning with our hypothesis, we observed that passive recoveries (e.g., dispersal mediated recolonization) elicited a lower magnitude of community response to a recovered predator than active recoveries (e.g., translocations, captive releases) (−0.45, 95% CI [−0.63, −0.27]) (Fig. 1, 3). We also found that systems which already supported one or more predator species elicited a higher magnitude of community response than systems without any existing predator (0.38, 95% CI [0.21,0.55]). Aside from recovery methods and existing community composition, no other covariate was supported (Table S2).

**Figure 3.**
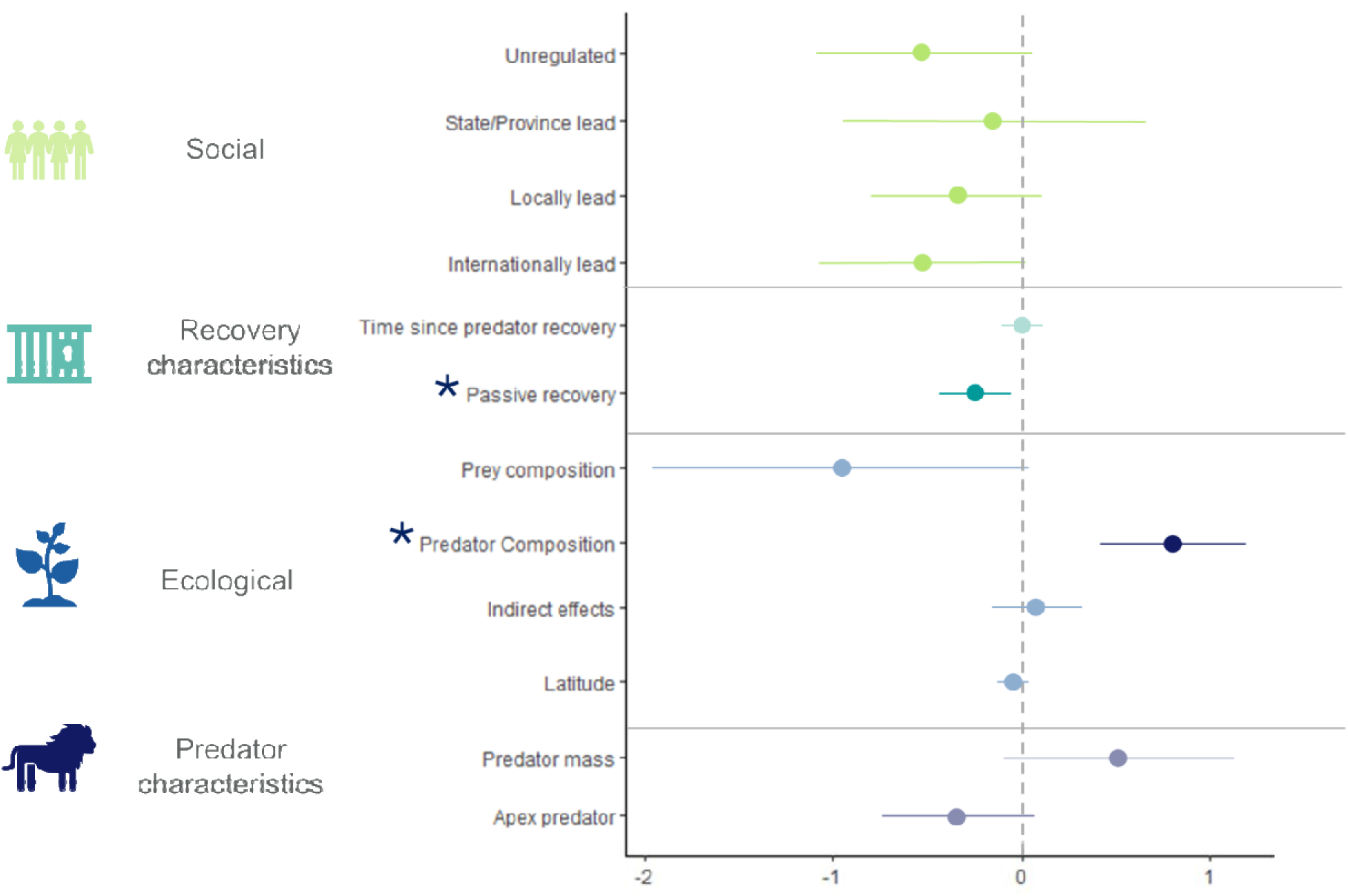
Coefficient plot and 95% CI for the global mixed effects model explaining the magnitude of community response to predator recovery. Saturated and starred predictors represent significance.

Community outcome alignment varied with the type of recovery method, the existing predator community composition, and the category of the community outcome (Fig. 3, Table S3). Like our previous linear model, we found that active recoveries were more likely to align with local conservation values and management preferences. We also found that systems with existing predators were less likely to have aligned outcomes. Finally, resulting behavioral community outcomes were less likely to align with local conservation values and management preferences, while changes in biogeochemical cycling and life history were more likely to align.

## Discussion

Predator recovery is often presumed to restore ecological functions and structure to biological communities, but the generality of this hypothesis, which undergirds many rewilding initiatives, is rarely systematically investigated. Here, we provide novel evidence that the downstream ecological outcomes of predator recovery are significantly impacted by existing predator community composition and mode of recovery (active or passive). Our comprehensive meta-analysis for the first time demonstrates that, while ecological effects of predator recovery are not ubiquitous, active predator reintroductions elicited stronger community responses than passive population recovery. More diverse extant predator communities were also significantly associated with greater effect magnitude following predator recovery.

The majority of community outcomes of predator recovery were aligned or neutral with local conservation values, with very few cases being unaligned (Fig. 4). The strong pattern of predator recovery producing positive conservation outcomes suggests that restoring predators can bolster ecosystem health without widespread unforeseen negative consequences. However, our meta-analysis is limited to reported community outcomes, and some negative community responses may be unreported in the literature. Similarly, human community outcomes are vastly unreported in conservation literature (Serota et al. 2023). Indeed, studies that focused solely on the ecological outcomes of predator reintroductions may have neglected to investigate recovered predator impacts on human communities, despite the fact that recovery projects that consider social dimensions have a greater likelihood of success (Serota et al. 2023).

**Figure 4.**
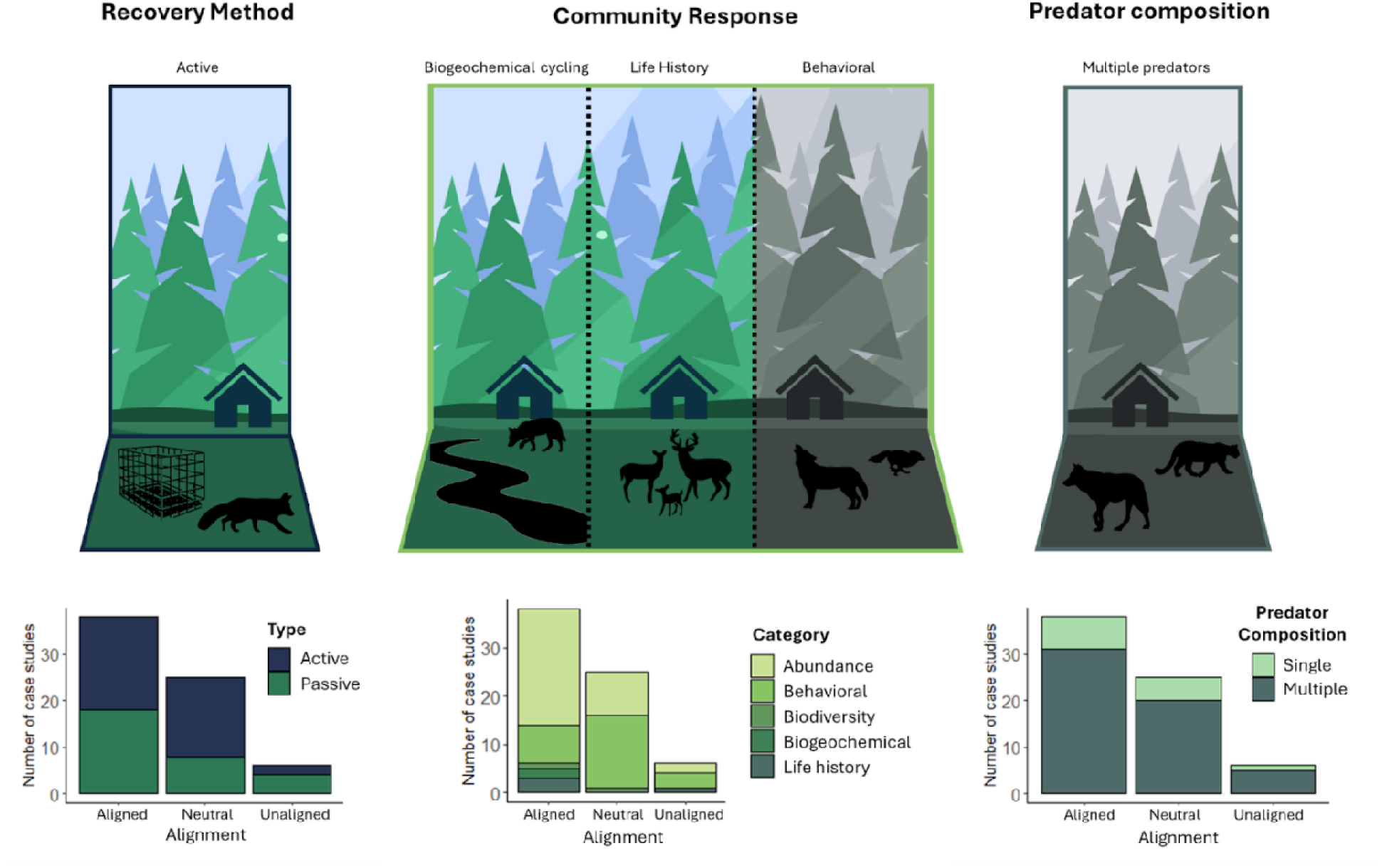
Graphical depiction of the significant factors driving outcome alignment. Panels in color represent positive coefficients, while panels in grayscale represent negative coefficients. Below are the distributions of recovery method, community outcomes, and existing predator composition at each recovery site, partitioned by alignment to local conservation values and management preferences.

We observed that passively recovered predators elicited a lower magnitude of community response and were less likely to have results aligned with local conservation values and management preferences. Often in passive recovery, the initial stressors that caused the extirpation are still present, meaning the predator is simply re-exposed (Palmer et al. 2010). These sentiments especially hold true if there are anti-predator concerns from adjacent human populations that go unaddressed, such as the potential for carnivore-livestock conflict without necessary mitigation or management (O’Rourke 2014). However, in active restoration, these stressors are often removed or considered before the reintroduction, often with the involvement of local governing bodies that may be more familiar with the threats that recovered predators might face (Serota et al. 2023). This does not mean that passive recoveries do not have substantial community outcomes, as 48% of the community outcomes with significant effects were from passive recoveries. Alternatively, our results may suggest that rather than active restoration being a more effective recovery tool objectively, intentional passive restoration (by means of connectivity efforts or habitat conservation and restoration) may also benefit from anticipation of local conditions faced by dispersing predators (Serota et al. 2023). For example, with legal protection under the Irish Wildlife Act (1976), European pine martens (*Martes martes)* passively recovered into Ireland, where they were able to cause significant changes to invasive gray squirrel *(Sciurus carolinensis)* activity (National Parks and Wildlife Service 1976, Twining et al. 2021). Similarly, community responses to passive predator recovery may only appear after a time-lag compared to community responses to active recovery, as passive recoveries themselves occur gradually while active recoveries consist of a short, immediate period of predator introduction (Hayward and Somers 2009). Many passively recovered predator populations have recovered fairly recently, and thus may lack the length of establishment time needed to precipitate measurable community responses (Watts et al. 2020).

We also found that the existing predator community composition positively influenced the magnitude of the community response, yet prey community composition was not a strong predictor of community outcome. In contrast, predator recovery in a multi-predator system was less likely to align with local management and conservation preferences (Fig. 5, Box 1). Our results counter classic interaction strength theory, which predicts that species in simpler communities are more likely to exert strong interactions on each other (Kokkoris et al. 2002, Berlow et al. 2004). While our search yielded a larger number of multi-predator systems, the observed positive effect could also be because a recovering predator not only adds functional redundancy but also can compound existing functional predator roles (Selkoe et al. 2015, Stier et al. 2016). Recovered predators also can have multiple interaction pathways in more complex systems, with community outcomes that have the potential to misalign with conservation values (Stepkovitch et al. 2023)

**Figure 5.**
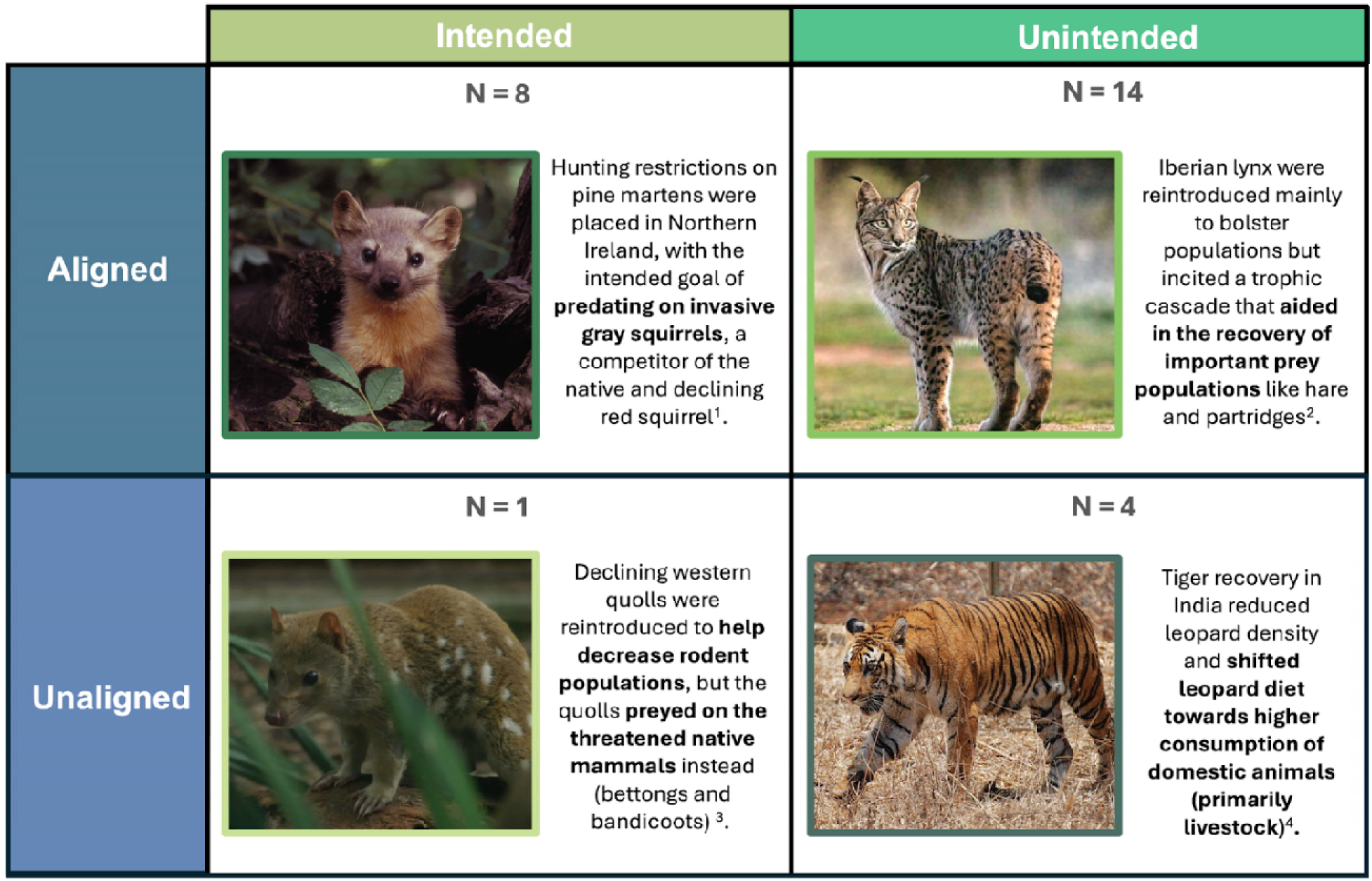
Examples of predator recovery intentions and whether they aligned with local conservation values. 1. Twining et al. 2021; 2. Jiménez et al. 2019, 3.Stepkovitch et al. 2023, 4. Harihar et al. 2011.

Behavioral community outcomes were less likely to align with conservation values and preferences, while community outcomes that altered biogeochemical cycling or life history traits were more likely to align. When a dominant predator recovers in an area, the recovered population has the potential to shift lower-trophic-level community interactions via both abundance and behavior. While behavioral shifts between predator and prey are commonly studied in reintroductions, we observed that behavioral responses within a carnivore guild were more common. Indeed, of all community outcomes with a resulting behavioral shift, 80% reported a shift in a subordinate predator, and only 20% reported a shift in prey behavior (Fig. 4). In subordinate carnivores, behavioral changes may manifest as prey-switching to a more accessible prey, like livestock. For instance, in areas where leopards (*Panthera pardus)* were sympatric with recovering tigers (*Panthera tigris)*, the subordinate leopard shifted its diet to consume more domestic prey (Harihar et al. 2011). Alternatively, altered biogeochemical processes or changes in life history traits may align more with local and community values. These responses are emblematic of the ecosystem services that predators provide humans, and might be looked upon favorably as they may be viewed as restoring an element of historic “balance” to an ecosystem (Wallach et al. 2015, Lennox et al. 2022). However, because these community responses require additional resources and time to monitor, indirect responses may be underreported in the literature. Furthermore, the foresight to measure and monitor lower trophic levels is often predicated on the expectation that they will change post-predator recovery.

While our results indicate that community responses to recovered predators are common, we note that a vast majority of predator recovery projects only reported metrics of the recovering predator population (e.g. survival, reproduction, etc.). Information on community outcomes may exist in gray literature or organizational project management plans, but these data were substantially lacking in published literature. However, we acknowledge that community responses are often harder to monitor. Subsequent responses in the community can be diverse, and indirect effects or trophic cascades may only be observable after a time lag, if they are observable at all (Peckarsky et al. 2008). Furthermore, passively recovering species are also more likely to go unnoticed and unmonitored. Additional insights on the influence of predator population recovery on ecological communities would be gained with improved study design before, during, and after predator restoration, and increased monitoring and reporting of direct and indirect effects in recovery zones.

This gap in knowledge on community outcomes highlights the importance of active management and monitoring in predator reintroductions. Only 9% of variation in community response strength was explained by our best model, indicating that socioecological complexity and context-dependency may make predicting the ecological role of recovered predators challenging in new scenarios. In addition, while a vast majority of community outcomes were aligned with local conservation and management values, the few that did not align illuminate the importance of consistent monitoring of both the predator population and the surrounding community. The predator impacts that ripple into the adjacent community are the most likely to interact with human presence, and can span multiple diverse pathways, such as beneficial ecosystem services (e.g. the regulation of overabundant mesopredators) or adverse human-wildlife conflicts (e.g. depredation events; (Harihar et al. 2011, Jiménez et al. 2019). Our analysis highlights the need for further interdisciplinary research between project managers, researchers, and local communities working in partnership towards a shared goal of conserving and restoring functional ecosystems (Serota et al. 2023).

Our results further bring into question how to best realize the intentions behind predator recoveries. Predators are unique in that they provide necessary ecosystem services and strong top-down interactions on lower trophic levels, while paradoxically, it is these strong top-down forces that drive adverse interactions between predators and humans (Nyhus 2016, Lennox et al. 2022). The same predator behaviors can simultaneously be viewed as positive or negative depending on the predator’s proximity to and relationship with humans. However, restoration rarely occurs in a vacuum, and considering the large home range requirements of many predators, expansion beyond the initial reintroduction site is likely, given enough time (Berger-TAL and Saltz 2014, Buderman et al. 2018, Hill et al. 2019) Accounting for the rapid expansion of socioecological landscapes, future predator recoveries must consider the probability of increased human-wildlife interactions, with recognition that the recovered predator’s behavior largely remains constant, and instead it is our values and preferences that are context dependent.

## Notes

### Competing Interest Statement

The authors have declared no competing interest.

