## Supplemental Figures for "Functional Roles of the World’s Recovered Predators"

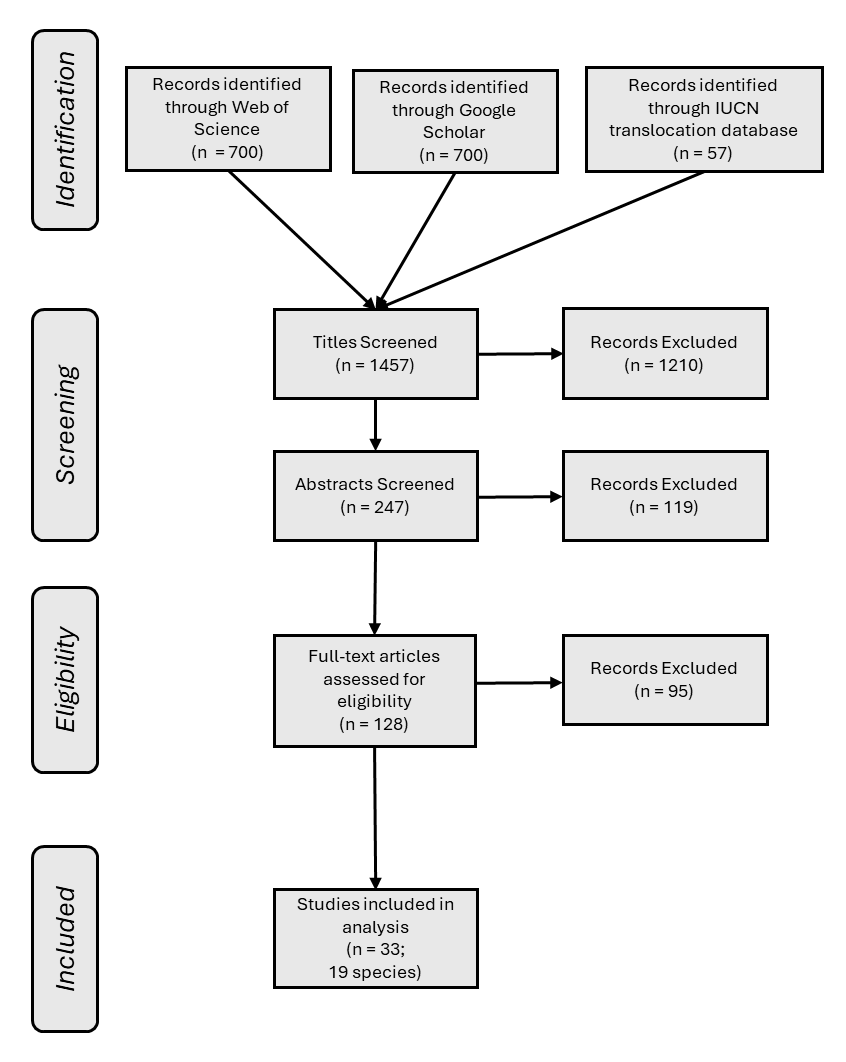

Figure S1. PRISMA  (Preferred Reporting Items for Systematic Reviews and Meta-Analyses) diagram indicating the selection criteria for recording community outcomes (Moher et al. 2015)

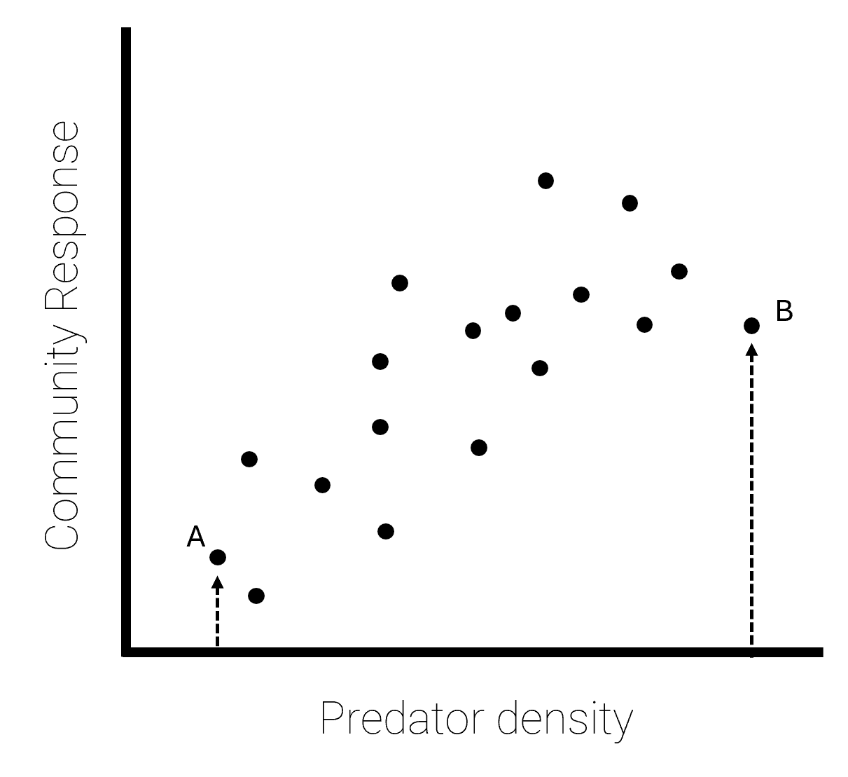

Figure S2.  Theoretical illustration of which community responses were selected (A and B) if a manuscript reported a gradient of predator density.

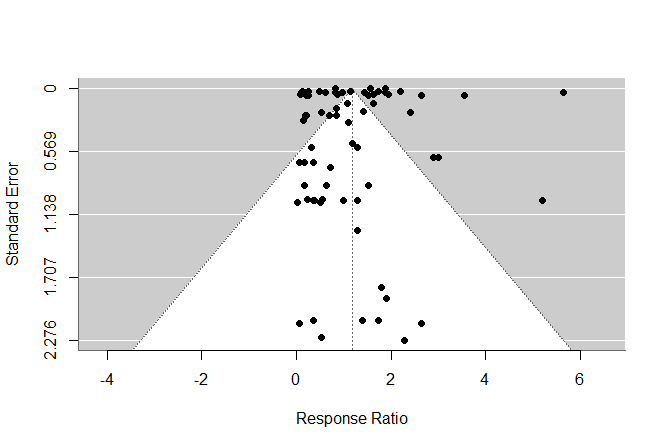

Figure S3. Funnel plot depicting the relationship between community response (Response Ratio), and standard error.

| Predictor | Reasoning | Citation |
| --- | --- | --- |
| Time since the predator was last extant in the area | We hypothesized that predators with a shorter duration of absence would elicit a higher magnitude of community response. | (Eitam and Blaustein 2010) |
| Whether the predator was an apex or mesopredator | We hypothesized that apex predators would elicit a larger magnitude of community response given their ability to affect multiple lower trophic levels. | (Ripple et al. 2014) |
| Latitude of predator recovery | We hypothesized that sites closer to the equator would have a larger vegetative community, which might provide a more complex system for the predator to affect, thus increasing the magnitude of response. | (Willig et al. 2003) |
| Whether the predator elicited a direct or indirect response | We hypothesized that predators that elicited a trophic cascade would elicit a larger magnitude of community response, as they affect a larger proportion of the community. | (Estes et al. 2011) |
| Mass of the predator | We hypothesized that larger predators would have a great effect on the community as they can exert strong top-down forces. | (Ripple et al. 2014) |
| Type of reintroduction (active or passive) | We hypothesized that active translocations would lead to a larger community response, as active translocation would have already accounted for ecological and anthropogenic factors that caused extirpation originally. | (Palmer et al. 2010) |
| Governing body that led the reintroduction | We hypothesized more locally led reintroductions would result in a larger community response, as locally driven translocations result in a higher rate of establishment. | (Serota et al. 2023) |
| GDP of the country where the reintroduction took place | We hypothesized that wealthier countries would have more resources to devote to maintaining and supporting a recovered predator. | (Naidoo and Adamowicz 2001) |
| Existing Predator community composition (single or multiple) in reintroduction area | We hypothesized that studies with a more complex predator community would cause a larger community response, as the predation pressure from multiple predators would compound onto lower trophic levels. | (Rosenfeld 2002) |
| Existing prey community composition (single or multiple) in reintroduction area | We hypothesized that predators introduced into sites with a more complex prey community would elicit a larger community response, as this would give the reintroduced predator multiple sources of food and thus functional impact. | (Garrott et al. 2007) |

Table S1. A list of fixed predictors included in the model competition explaining the magnitude of community response to a reintroduced predator

| Predictors | AICC | Delta AICC | Weights |
| --- | --- | --- | --- |
| Magnitude of community response ~ (1\| Study) + (1\| Species) + Type of Recovery + existing predator composition | 206.47 | 0 | 0.077 |
| Magnitude of community response ~ (1\| Study) + (1\| Species)  + Latitude + Type of Recovery + existing predator composition | 206.80 | 0.33 | 0.065 |
| Magnitude of community response ~ (1\| Study) + (1\| Species)  + Latitude + predator mass + Type of Recovery | 208.46 | 1.99 | 0.029 |
| Magnitude of community response ~ (1\| Study) + (1\| Species)  + Years since extirpation + Type of Recovery + existing predator composition | 208.49 | 2.02 | 0.028 |
| Magnitude of community response ~ (1\| Study) + (1\| Species)  + indirect pathway + Type of Recovery + existing predator composition | 208.5 | 2.03 | 0.028 |

Table S2. Top five performing models demonstrating what factors influence the magnitude of community response to predator recovery.

| Predictors | AICC | Delta AICC | Weights |
| --- | --- | --- | --- |
| Alignment with Local conservation preferences ~ (1\| Study) + (1\| Species) + Type of Recovery + existing predator composition + category of community response | 55.1 | 0 | 0.890 |
| Alignment with Local conservation preferences ~ (1\| Study) + (1\| Species)  + GDP of country where recovery took place + Type of Recovery + existing predator composition | 60 | 0.33 | 0.076 |
| Alignment with Local conservation preferences ~ (1\| Study) + (1\| Species)  + GDP of country where recovery took place + existing predator composition + Type of Recovery + outcome intention | 63 | 1.99 | 0.017 |
| Alignment with Local conservation preferences ~ (1\| Study) + (1\| Species)  + outcome intention + predator mass + existing predator composition | 63.1 | 2.02 | 0.016 |
| Alignment with Local conservation preferences ~ (1\| Study) + (1\| Species)  + category of community response GDP of country where recovery took place + existing predator composition  + outcome intention | 69.4 | 2.03 | 0.001 |

Table S3. Top five performing models demonstrating what factors influence whether a community outcomes is aligned with local conservation and management preferences.
